# CASTLE: A database of synthetic lethal sets predicted from genome-scale metabolic networks

**DOI:** 10.1101/2021.02.08.430024

**Authors:** Vimaladhasan Senthamizhan, Sunanda Subramaniam, Arjun Raghavan, Karthik Raman

## Abstract

**Summary:** Genome-scale metabolic networks have been reconstructed for hundreds of organisms over the last two decades, with wide-ranging applications, including the identification of drug targets. Constraint-based approaches such as flux balance analysis have been effectively used to predict single and combinatorial drug targets in a variety of metabolic networks. We have previously developed Fast-SL, an efficient algorithm to rapidly enumerate all possible synthetic lethals from metabolic networks. Here, we introduce CASTLE, an online standalone database, which contains synthetic lethals predicted from the metabolic networks of over 130 organisms. These targets include single, double or triple lethal set of genes and reactions, and have been predicted using the Fast-SL algorithm. The workflow used for building CASTLE can be easily applied to other pathogenic models and used to identify novel therapeutic targets.

**Availability:** CASTLE is available at https://ramanlab.github.io/CASTLE/

## 1 Introduction

The recent years have witnessed the reconstruction of a number of genome-scale metabolic models [1, 2], including pathogenic organisms [3–5], with applications in drug target identification and understanding disease aetiology [6–10]. These genome-scale models of cellular metabolism capture all known reactions in an organism’s metabolic network, and are particularly useful to predict the growth rate of an organism, and its phenotype upon various perturbations, particularly the removal of one or more genes/reactions [11]. These networks have been studied using Flux Balance Analysis (FBA) and FBA has proven to accurately predict phenotypes following various genetic perturbations [12, 13]. FBA has been employed as a key strategy in antimicrobial discovery pipelines [9, 14]. FBA can also be used to reliably predict synthetic lethal genes in metabolic network which in turn can be used to identify combinatorial targets for various pathogens [15, 16]. These combinatorial targets, which are termed ‘Synthetic Lethal Sets’ are sets of reactions/genes, in which only the simultaneous deletion of all reactions/genes in the set will abrogate growth and lead to the subsequent death of the organism. (Figure 1).

**Figure 1:**
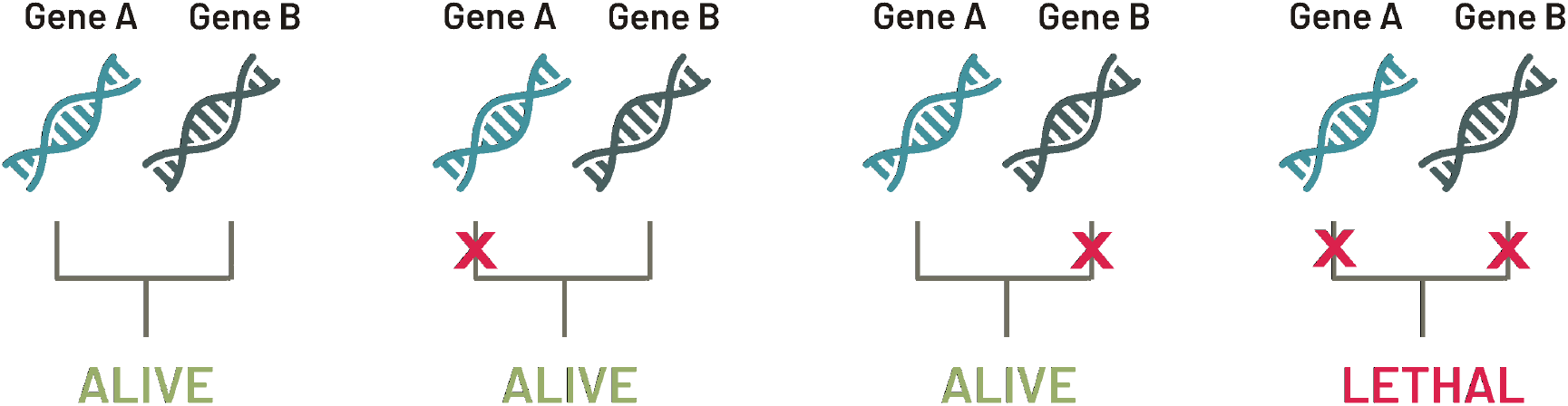
Schematic illustrating the logic of synthetic lethality. If there are two targets A and B, deleting any one of them does not affect the survival of the organism. Only the simultaneous deletion of both targets has a lethal effect on the organism. In this case, targets A and B are known as ‘double lethals’. This logic can be extended to ‘single lethals’ and ‘triple lethals’

Since the systematic evaluation of such combinatorial deletions in vivo is challenging, computational approaches have been of great interest to overcome this difficulty. A number of algorithms have been developed to predict synthetic lethals in metabolic networks [17–19]. Fast-SL, previously developed in our laboratory, is as yet the most efficient parallel algorithm available to predict synthetic lethal sets in metabolic networks [20]. Fast-SL algorithm circumvents the computational complexity of various other methods through an iterative reduction of the search space for higher order combinatorial targets. Many studies have used Fast-SL to predict synthetic lethals [21–24] and explore them as possible drug targets. To further enable the analysis of such combinatorial deletions across organisms, we used Fast-SL to computationally identify synthetic lethal sets for a variety of pathogenic organisms. The results of the analyses have been compiled and published in CASTLE (Computational Analysis of SynThetic LEthals) - a standalone web database that we report here.

## 2 Methods

In order to identify synthetic lethal sets of genome-scale metabolic models, we used the parallel version of the Fast-SL program. A MATLAB script was written to interface the COBRA Toolbox [11] with the parallel Fast- SL programs and automate the process of identifying single, double, and triple reaction/gene lethal sets for a group of pathogenic organisms. The metabolic model was first read using the COBRA toolbox; the parallel Fast-SL script was then employed to identify the lethals, which were written out as MAT files.

The genome-scale metabolic models themselves were selected and downloaded from the BiGG database [25] and Virtual Metabolic Human (VMH) database [26]. The output results on lethal information were saved as an Excel file in CSV format. These lethal sets were converted into three different formats (JSON, CSV, MAT) to offer users choice in the format they can view the data. This process was repeated for all the 113 organisms and the results were compiled to be published in the database. The names of all organisms along with the information on metabolic models and growth rate are given in Supplementary Information.

## 3 Results

The synthetic lethal sets (single, double and triple lethal reactions and genes) of 113 pathogenic organisms were identified. The distribution of the number of synthetic lethal sets identified by Fast-SL algorithm is shown in Figure 2. This information was incorporated into CASTLE—a standalone web database. In the home page, the number of each type of lethal set which can be selected to open further information about the reactions/genes. Further, a download icon is available, which, when selected, will automatically download all available results as a zip file.

**Figure 2:**
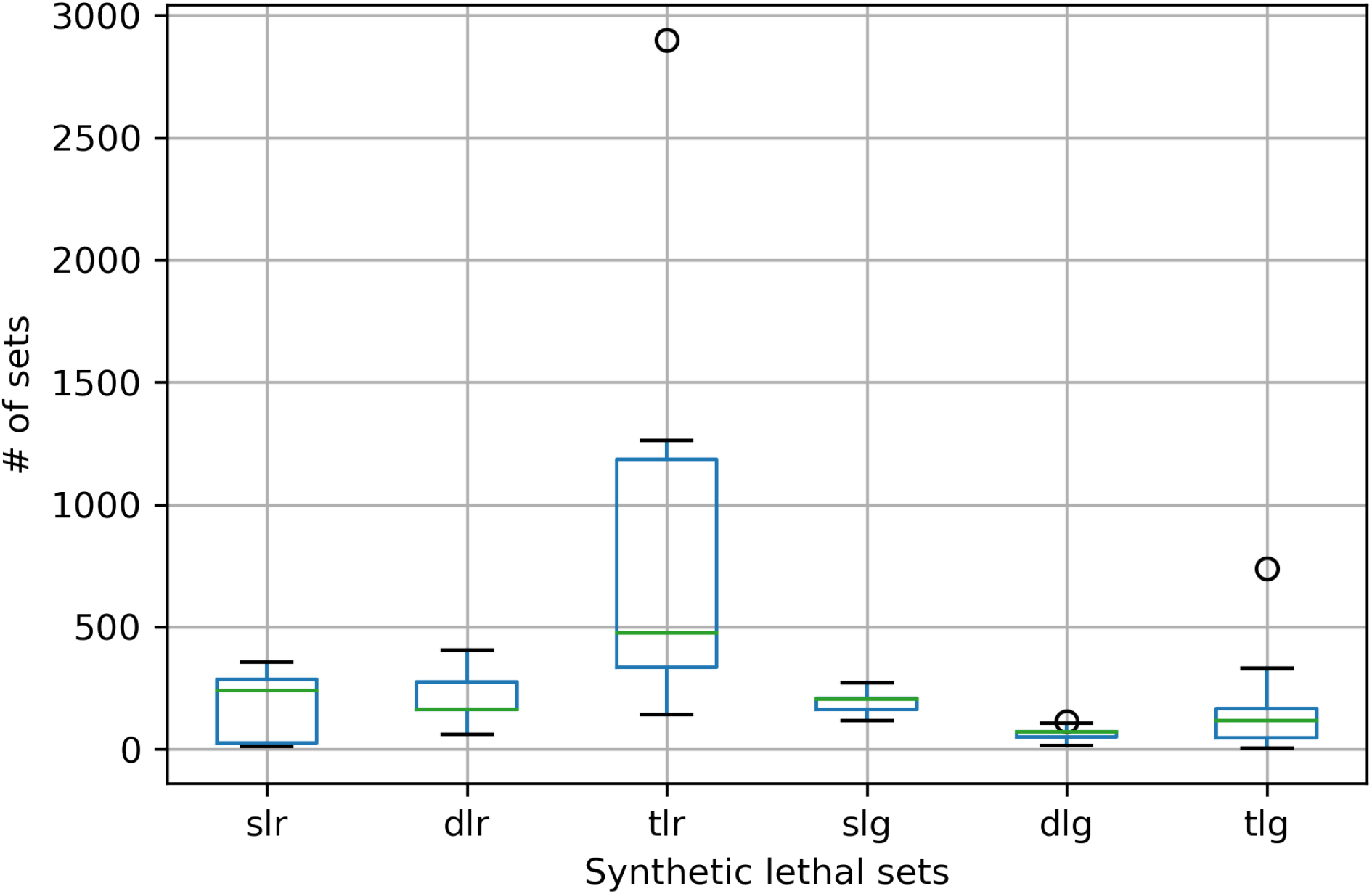
Box plot showing the distribution of the synthetic lethal sets identified in the 113 organisms. The larger box plot of ‘tlr’ is heavily influenced by the sizable cohorts of triple lethals sets from Escherichia coli and Shigella organisms. **Abbreviations:** slr - Single Lethal Reactions; dlr = Double Lethal Reactions; tlr = Triple Lethal Reactions; slg = Single Lethal Genes; dlg = Double Lethal Genes; tlg = Triple Lethal Genes

The database also features individual organism page which contains the list of synthetic lethal sets for that particular organism. Equipped with smooth-flowing anchors for better user experience, the individual organism page also provides you with hyperlinks for each synthetic lethal reaction/gene which takes the user to VMH and BiGG databases for further information about the target.

Finally, we provide our synthetic lethal sets as downloadable files available in the organism-specific page. A user can opt for any of the three file formats (JSON, CSV and MAT files) available. The ‘About’ page provides the user with a brief overview of synthetic lethals and contains instructions on how to use the database effectively.

## 4 Discussion

In this study, synthetic lethal sets (single, double, and triple) were found for 133 pathogenic organisms using the COBRA Toolbox and Parallel Fast-SL for MATLAB. This data was made accessible to global research community through an online database. This database is developed with a motive to fill an existing gap in this area of research, by providing the research community with an easy-to-access, standardized resource unlike any other currently in existence. This database is envisaged to help researchers tackle the twin problem of drug resistance and drug side effects mentioned previously, as well as discover new therapeutic targets. Furthermore, the standardized procedure used can be easily extended to the models of other pathogenic models, which have not been created yet / have not been analyzed in this study.

## Acknowledgments

VS acknowledges Initiative for Biological Systems Engineering (IBSE) for the post-baccalaureate fellowship. AR acknowledges funding from Mathematics of Information Technology and Complex Systems (MITACS) and support from Mr. Ryan Caldwell and others at MITACS.

